# *scater:* pre-processing, quality control, normalisation and visualisation of single-cell RNA-seq data in R

**DOI:** 10.1101/069633

**Authors:** Davis J. McCarthy, Kieran R. Campbell, Aaron T. L. Lun, Quin F. Wills

## Abstract

**Motivation:** Single-cell RNA sequencing (scRNA-seq) is increasingly used to study gene expression at the level of individual cells. However, preparing raw sequence data for further analysis is not a straightforward process. Biases, artifacts, and other sources of unwanted variation are present in the data, requiring substantial time and effort to be spent on pre-processing, quality control (QC) and normalisation.

**Results:** We have developed the R/Bioconductor package *scater* to facilitate rigorous pre-processing, quality control, normalisation and visualisation of scRNA-seq data. The package provides a convenient, flexible workflow to process raw sequencing reads into a high-quality expression dataset ready for downstream analysis. *scater* provides a rich suite of plotting tools for single-cell data and a flexible data structure that is compatible with existing tools and can be used as infrastructure for future software development.

**Availability:** The open-source code, along with installation instructions, vignettes and case studies, is available through Bioconductor at http://bioconductor.org/packages/scater.

**Supplementary information:** Supplementary material is available online at *bioRxiv* accompanying this manuscript, and all materials required to reproduce the results presented in this paper are available at dx.doi.org/10.5281/zenodo.60139.

## Introduction

Single-cell RNA sequencing (scRNA-seq) describes a broad class of techniques which profile the transcriptomes of individual cells. This provides insights into cellular processes at a resolution that cannot be matched by bulk RNA-seq experiments (Hebenstreit and Teichmann, 2011; Shalek et al., 2013). With scRNA-seq data, the contributions of different cell types to the expression profile of a heterogeneous population can be explicitly determined. Rare cell types can be interrogated and new cell subpopulations can be discovered. Graduated processes such as development and differentiation can also be studied in greater detail. However, this improvement in resolution comes at the cost of increased technical noise and biases. This means that pre-processing, quality control and normalisation are critical to a rigorous analysis of scRNA-seq data. The increased complexity of the data across hundreds or thousands of cells also requires sophisticated visualisation tools to assist interpretation of the results.

Numerous statistical methods and software tools have been published for scRNA-seq data (Guo et al., 2015; Kharchenko et al., 2014; Finak et al., 2015; Angerer et al., 2015; Juliá et al., 2015; Trapnell et al., 2014). However, all of these assume that quality control and normalisation have already been applied. Fewer methods are available in the literature to perform these basic steps in scRNA-seq data processing (Ilicic et al., 2016). This issue is exacerbated by the diversity of scRNA-seq data sets with respect to the experimental protocol and the biological context of the study, meaning that a single processing pipeline with fixed parameters is unlikely to be universally applicable. Rather, software tools are required that support an interactive approach to analysis. This allows parameters to be fine-tuned for the study at hand in response to any issues diagnosed during data exploration. The provided functionality should also process the data in a statistically rigorous manner and encourage reproducible bioinformatics analyses.

One of the most popular frameworks for interactive analysis is the R programming language, extended for biological data analysis through the Bioconductor project (Huber et al., 2015). While Bioconductor packages have been widely used for bulk RNA-seq data, the existing data structures (like the ExpressionSet class) are not sufficient for scRNA-seq data. This is because they do not support data types that are specific to single-cell studies, e.g., cell-cell distance matrices for clustering. For larger studies, this also includes data beyond expression profiles such as intensity values from fluorescence-activated cell sorting, cell imaging data, and information from epigenetic and targeted genotyping assays. Existing methods for processing and applying quality control to scRNA-seq data are similarly inadequate. In particular, current visualisation methods designed for exploratory data analysis of bulk transcriptomic experiments are unsuited to datasets containing hundreds or thousands of cells. The large size of each dataset also favours methods such as *kallisto* (Bray et al., 2016) and *Salmon* (Patro et al., 2015) for rapidly quantifying gene expression. Extensions to the current computational infrastructure are required to provide appropriate data structures and methods that can accommodate these rich scRNA-seq datasets for integrative analyses of expression and other assay data along with the accompanying metadata.

Here we present *scater*, an open-source R/Bioconductor software package that implements a convenient data structure for representing scRNA-seq data and contains functions for pre-processing, quality control, normalisation and visualisation. The package provides wrapper functions for running *kallisto* and *Salmon* on raw read data and converting their output into gene-level expression values, methods for computing and visualising quality-control metrics for cells and genes, and methods for normalisation and correction of uninteresting covariates. This is done in a single software environment which enables seamless integration with a large number of existing tools for scRNA-seq data analysis in R. The *scater* package provides basic infrastructure upon which customized scRNA-seq analyses can be constructed, and we anticipate the package to be useful across the whole spectrum of users, from experimentalists to computational scientists.

## Methods, Data and Implementation

### Case study with scRNA-seq data

The results presented in the main paper and supplementary case study use an unpublished single-cell RNA-seq dataset consisting of 73 cells from two lymphoblast cell lines of two unrelated individuals. Cells were captured, lysed, and cDNA generated using the popular C1 platform from Fluidigm, Inc. (www.fluidigm.com/products/c1-system). The processing of the two cell lines was replicated across two machines, with the nuclei of the two cell lines stained with different dyes before mixing on each machine. Cells were imaged before lysis, with an example image provided together with these data (see Case Study in Supplementary Material). Samples were sequenced with paired-end sequencing using the HiSeq 2500 Sequencing system (Illumina). RNA-seq reads were were mapped to a custom genome reference, consisting of Homo sapiens GRCh37 (primary assembly from ftp.ensembl.org/pub/release-75/fasta/homo_sapiens/dna/, last accessed 14.08.2015), Epstein-Barr Virus type 1 (B95-8 strain, Accession NC 007605.1) and ERCC RNA spike-ins (ThermoFisher). Reads in fastq format were aligned using TopHat2 v2.0.12 (Kim et al., 2013) using Bowtie2 v2.2.3.0 (Langmead and Salzberg, 2012) as the core-mapping engine (`--mate-inner-dist 190 --mate-std-dev 40 --report-secondary-alignments`) and other default parameters. Potential PCR duplicates were marked with Picard MarkDuplicates v1.92(1464). Reads mapping uniquely to annotated exon features were counted using htseq-count implemented in HTSeq, version 0.6.1p1 (Anders et al., 2015).

Further case studies using *scater* on published data, for example from 3000 mouse cortex cells (Zeisel et al., 2015) and 1200 cells from early-development mouse embryos (Scialdone et al., 2016) are available at dx.doi.org/10.5281/zenodo.59897. All materials required to reproduce the results presented in this paper are available at dx.doi.org/10.5281/zenodo.60139.

### Implementation

The *scater* package is an open-source R package available through Bioconductor. Key aspects of the code are written in C++ to minimise computational time and memory use. The package builds on many other R packages, including *Biobase* and *BiocGenerics* for core Bioconductor functionality (Huber et al., 2015); *destiny* (Angerer et al., 2015) and *Rtsne* for dimensionality reduction; and *edgeR* (Robinson et al., 2010) and *limma* (Ritchie et al., 2015) for model fitting and statistical analyses. The plotting functionality in the package uses *ggplot2* (Wickham, 2016). A full set of dependencies is provided in the Supplementary Materials.

## Results

### The *scater* package

The *scater* package offers a workflow to convert raw read sequences into a data set ready for higher-level analysis within the R programming environment (Figure 1). In addition, *scater* provides basic computational infrastructure to standardise and streamline scRNA-seq data analyses. Key features of *scater* include: (1) the “single-cell expression set” (SCESet) class, a data structure specialized for scRNA-seq data; (2) wrapper methods to run *kallisto* and *Salmon* and process their output into gene-level expression values; (3) automated calculation of quality control metrics, with QC visualisation and filtering methods to retain high-quality cells and informative features; (4) extensive visualisation capabilities for inspection of scRNA-seq data; and (5) methods to identify and remove uninteresting covariates affecting expression across cells. The package integrates many commonly used tools for scRNA-seq data analysis and provides a foundation on which future methods can be built. The methods in *scater* are agnostic to the form of the input data and are compatible with counts, transcripts-per-million, counts-per-million, FPKM or any other appropriate transformation of the expression values.

**Figure 1:**
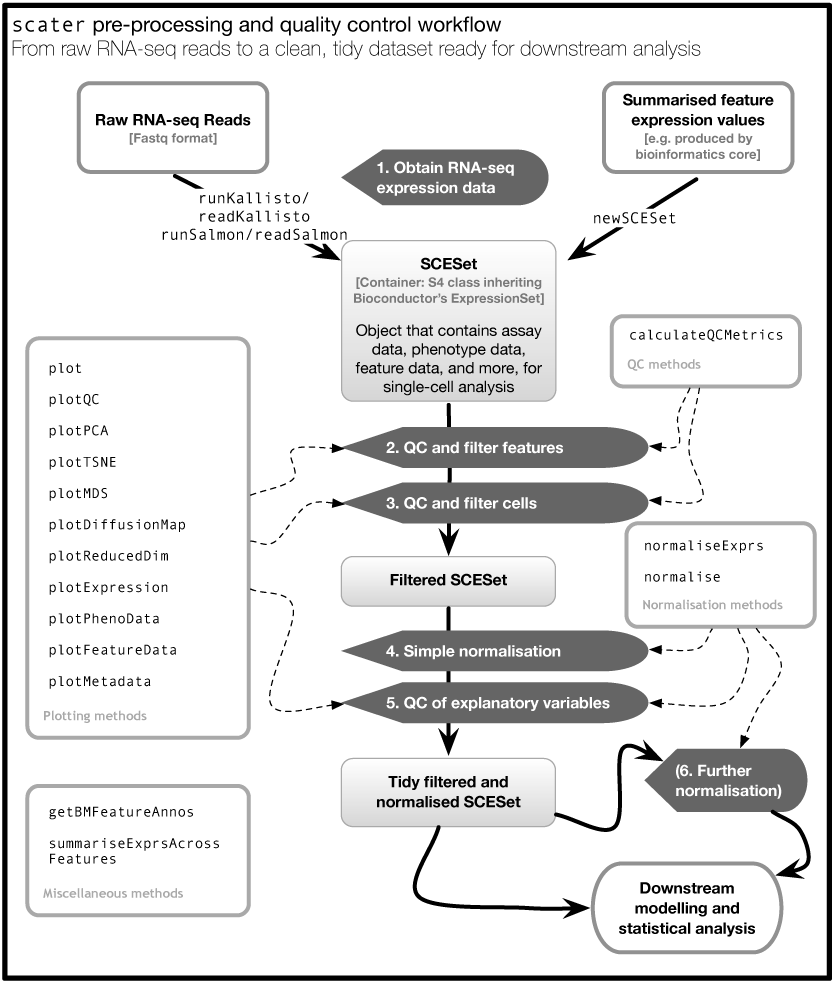
An overview of the *scater* workflow, from raw sequenced reads to a tidy data set ready for higher-level downstream analysis. For step 5, explanatory variables include experimental covariates like batch, cell source and other recorded information, as well as QC metrics computed from the data. Step 6 describes an optional round of normalisation to remove effects of particular explanatory variables from the data. Automated computation of QC metrics and extensive plotting functionality support the workflow.

### SCESet: a data structure for single-cell expression data

The *scater* package is built around the SCESet class (Supplementary Figure 1) which provides a sophisticated container for scRNA-seq data. This class inherits from the ExpressionSet class in Bioconductor’s *Biobase* package (Huber et al., 2015), which allows assay data (and multiple transformations thereof), gene or transcript metadata and sample metadata to be combined in a single object to empower robust analyses. While the ExpressionSet class is the basis of many microarray and bulk RNA-seq anaysis methods in Bioconductor, extensions to the class design are necessary to support scRNA-seq data analyses. Specifically, the SCESet class adds slots to store a reduced-dimension representation of the expression profiles, to easily visualize the relationships between cells; cell-cell and gene-gene pairwise distance matrices, for clustering or regulatory network reconstruction; bootstrapped expression results (such as from *kallisto*), to gauge the accuracy of expression quantification; consensus clustering results, where cluster assignments for each cell are combined from different methods to improve reliability; information about feature controls (such as ERCC spike-ins), which is required in downstream steps such as normalization, QC and detection of highly variable genes; and several more (Supplementary Figure 1). With these extra slots, SCESet objects can support analyses of scRNA-seq data that ExpressionSet cannot. In addition, extra data types such as FACS marker expression or epigenetic information can be easily stored in each SCESet object for integration with the single-cell expression profiles.

An SCESet data object can be easily subsetted by row or column to remove unwanted genes or cells, respectively, from all data and metadata fields stored in the object. Furthermore, data and metadata in multiple SCESet objects can be easily combined e.g., to incorporate cells from different experimental batches. SCESet objects can also be converted to other R data structures, or saved to disk in structured, shareable formats. Further details on the class, including its motivation and execution, are available in the Supplementary Case Study and the package documentation. All methods available in *scater* are applicable to instances of the SCESet class and exploit the availability and richness of (meta)data stored in each SCESet object.

### Data pre-processing

An important initial step in scRNA-seq data processing is to quantify the expression level of genomic features such as transcripts or genes from the raw sequencing data. Approaches to expression quantification from raw reads are, in principle, the same for scRNA-seq as they are for bulk RNA-seq (Kanitz et al., 2015; Teng et al., 2016). Read counts obtained from conventional quantification methods such as *HTSeq* (Anders et al., 2015) and *featureCounts* (Liao et al., 2014) can be readily stored in an SCESet object and used in a *scater* workflow (Figure 1). Another option is to use computationally-efficient pseudoalignment methods such as *kallisto* and *Salmon*. This is especially appealing for large scRNA-seq data sets containing hundreds to tens of thousands of cells. To this end, *scater* also provides wrapper functions for *kallisto* and *Salmon* so that fast quantification of transcript-level expression can be managed completely within an R programming environment. A common subsequent step for these methods is to collapse transcript-level expression to gene-level expression. Exploiting the *biomaRt* R/Bioconductor package, *scater* provides a convenient function for using Ensembl annotations to obtain gene-level expression values and gene or transcript annotations (Yates et al., 2016).

### Data quality control

The *scater* package provides methods to compute relevant QC metrics for an SCESet object. Given a set of control genes and/or cells, a variety of QC metrics will be computed and returned to the object in a single call to the calculateQCMetrics function (see package documentation). Cell-specific QC metrics include the total count across all genes, the total number of expressed genes, and the percentage of counts allocated to control genes like spike-in transcripts or mitochondrial genes. These metrics are useful for identifying low-quality cells—for example, a high percentage of counts mapping to spike-ins typically indicates that a small amount of RNA was captured for the cell, suggesting protocol failure or death of the cell in processing that renders it unsuitable for downstream analyses. For each gene, QC metrics such as the average expression level and the proportion of cells in which the gene is expressed are computed. This can be used to identify low-abundance genes or genes with high dropout rates that should be filtered out prior to downstream analyses. All of these metrics are used by scater to construct QC plots to diagnose potential issues with data quality. This facilitates quality control which—despite attempts at automation (Ilicic et al., 2016)—still requires manual intervention to account for aspects of the data specific to each study. The package documentation provides full details of the QC metrics produced.

In *scater*, the default plot method for an SCESet object produces a cumulative expression plot (Figure 2a). This plot describes how reads are distributed across genes, distinguishing between low-complexity libraries (where very few genes contain most of the counts) and their high-complexity counterparts (where counts are distributed more evenly across genes). For example, there is substantial variability in library complexity among cells in the case study dataset in Figure 2a. Some cells have profiles similar to the blank wells, suggesting that library preparation or sequencing failed for these cells and that the corresponding libraries should be removed prior to further analysis. Cell phenotype variables can be incorporated into these plots to highlight differences in expression distributions for different types of cells. For example, the curve for each cell is coloured by the type of well that produced the library (Figure 2a), while cells can also be split into separate facets by library type to show more metadata variables simultaneously (see Supplementary Case Study). Cumulative expression plots should be favoured over boxplots as the default method for visualising expression distributions across cells in a dataset, as the latter performs poorly at handling the long tail of low- and zero-expression observations in scRNA-seq data.

**Figure 2:**
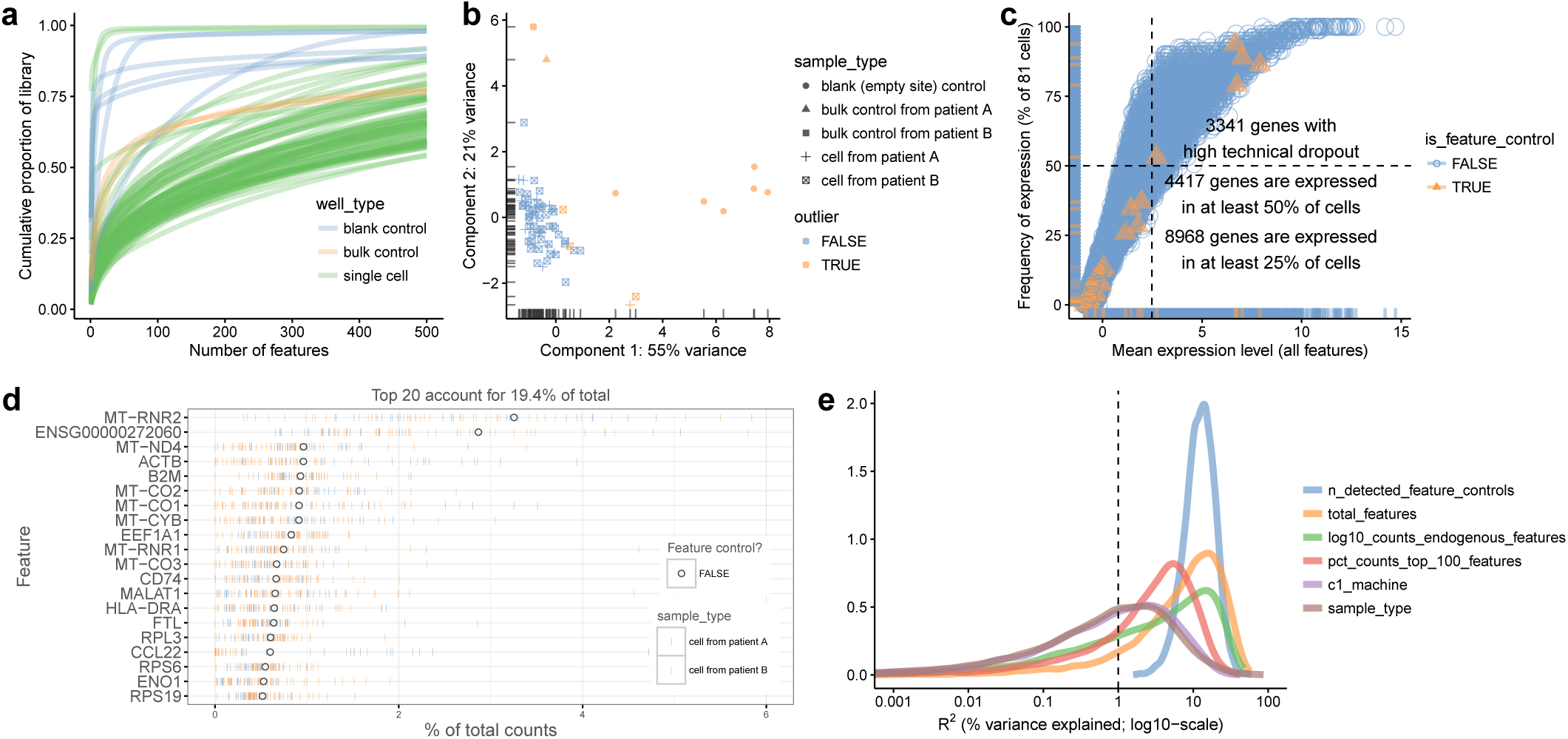
Different types of QC plots that can be generated with *scater*. (**a**) Cumulative expression plot showing the proportion of the library accounted for by the top 1–500 most highly-expressed features. (**b**) PCA plot produced using a subset of the QC metrics computed with *scater’s* calculateQCMetrics function. (**c**) Plot of frequency of expression (percentage of cells in which the feature is deemed expressed) against mean expression level across cells. The vertical dotted line shows the median of the gene mean expression levels, and the horizontal dotted line indicates 50% frequency of expression. (**d**) Plot of the 20 most highly-expressed features (computed according to the highest total read counts) across all cells in the data set. For each feature, the circle represents the percentage of counts that the gene accounts for computed from counts pooled across all cells. The genes are ordered by this value. The bars for each gene show the percentage of counts accounted for by the gene for each individual cell, providing a visualisation of the distribution across cells. (**e**) Density plot showing the percentage of variance explained by a set of explanatory variables across all genes. Each individual plot is produced by a single call with either the function plot (**a**), plotPCA (**b**) or plotQC (**c–e**).

The plotPCA function implements an approach to automatic outlier detection using multivariate normal methods applied to the cell-level QC metrics (Ilicic et al., 2016). Specifically, PCA is applied to the QC metrics for all cells and a plot is produced to automatically detect outliers in the higher-dimensional QC metric space (Figure 2b). These outliers correspond to low-quality cells with abnormal library characteristics (e.g., low total counts and few expressed genes) that should be removed prior to downstream analysis. This automated approach is powerful but also somewhat opaque with respect to how outliers are defined, and so complements simpler filtering approaches that apply thresholds to particular QC metrics.

The plotQC function generates many types of plots useful for quality control, such as a plot to visualise the frequency of expression of features against their average expression level (Figure 2c). Such plots are useful, because scRNA-seq is characterised by a high frequency of “dropout” events, that is, no observed expression (such as no read counts) in a particular cell for a gene that is actually expressed in that cell. Typically only a small set of genes are observed with detectable expression in every cell. With plotQC, control features can be highlighted easily in the plot, and typical scRNA-seq datasets will show a broadly sigmoidal relationship between average expression level and frequency of expression across cells. This is consistent with expected behaviour where genes with greater average expression are more readily captured during library preparation and are detected at a greater frequency (Brennecke et al., 2013; Kim et al., 2015; Vallejos et al., 2015).

With plotQC we can also produce a plot to visualise the most highly-expressed features in the dataset (Figure 2d). This provides a feature-centric overview of the dataset that visualises the features with highest total expression across all cells, while also displaying the distribution of cell-level expression values for these features. It is common to see ERCC spike-ins (if used), mitochondrial and ribosomal genes among the highest expressed genes, while datasets consisting of healthy cells will also show high levels of constitutively expressed genes like *ACTB*. This plot allows the analyst to quickly check that the gene- or transcript-level quantification is behaving as expected, and to flag datasets where it is not.

Another important step in quality control is to identify variables (experimental factors or computed QC metrics) that drive variation in expression data across cells. The plotQC function provides a novel approach to identifying variables that have substantial explanatory power for many genes. For each variable in the phenoData slot of the SCESet object, we fit a linear model for each feature with just that variable as the explanatory variable. We then plot the distribution of the marginal *R*^2^ values across all features for the variables with the most explanatory power for the dataset (Figure 2e). The variables are ranked by median *R*^2^ across features in the plot, allowing users to identify variables that may need to be considered during normalisation or statistical modelling. The plotQC function can also assess the influence of variables of interest by plotting principal components of the expression matrix most strongly correlated with a variable of interest against that variable. For example, in the Case Study data, the first principal component is correlated with the C1 machine used to process the cell (Supplementary Figure 2).

We also introduce the plotPhenoData function for convenient plotting of cell phenotype information (including QC metrics), and the plotFeatureData function for plotting feature information (see examples in the Supplementary Case Study). These methods will work not only on the SCESet class defined in *scater*, but also on any ExpressionSet object, providing sophisticated plotting functionality for many other Bioconductor packages and contexts.

The *scater* graphical user interface (GUI) provides convenient access to *scater*'s QC and visualisation methods (Supplementary Figures 4–6). This opens an interactive interface in a web browser that facilitates exploration of the data through QC plots and other intuitive visualisations. The GUI allows users of any background to easily examine the effects of changing multiple parameters, which can be helpful for quickly conducting exploratory data analysis. Useful settings can then be stored in R scripts to ensure that data analyses are reproducible.

In summary, *scater* provides a variety of novel and convenient methods to visualise an scRNA-seq dataset for QC. Low-quality cells and uninteresting genes can then be easily removed by filtering and subsetting the SCESet data structure prior to further analysis.

### Data visualisation

Dimensionality reduction techniques are necessary to convert high-dimensional expression data into low-dimensional representations for intuitive visualisation of the relationships, similarities and differences between cells. To this end, *scater* provides convenient functions to apply a variety of dimensionality reduction procedures to the cells in an SCESet object. Functions include plotPCA, to perform a principal components analysis; plotTSNE, to perform t-distributed stochastic neighbour embedding (Van der Maaten and Hinton, 2008), which has been widely used for scRNA-seq data (Amir et al., 2013; Bendall et al., 2014; Macosko et al., 2015); plotDiffusionMap, to generate a diffusion map (Haghverdi et al., 2015) for visualising differentiation processes; and plotMDS, to generate multi-dimensional scaling plots (Figure 3a–c). The plotReducedDim function can also be used to plot any reduced-dimension representation of cells (e.g., an independent component analysis produced by *monocle* (Trapnell et al., 2013) or similar) that is stored in an SCESet object.

**Figure 3:**
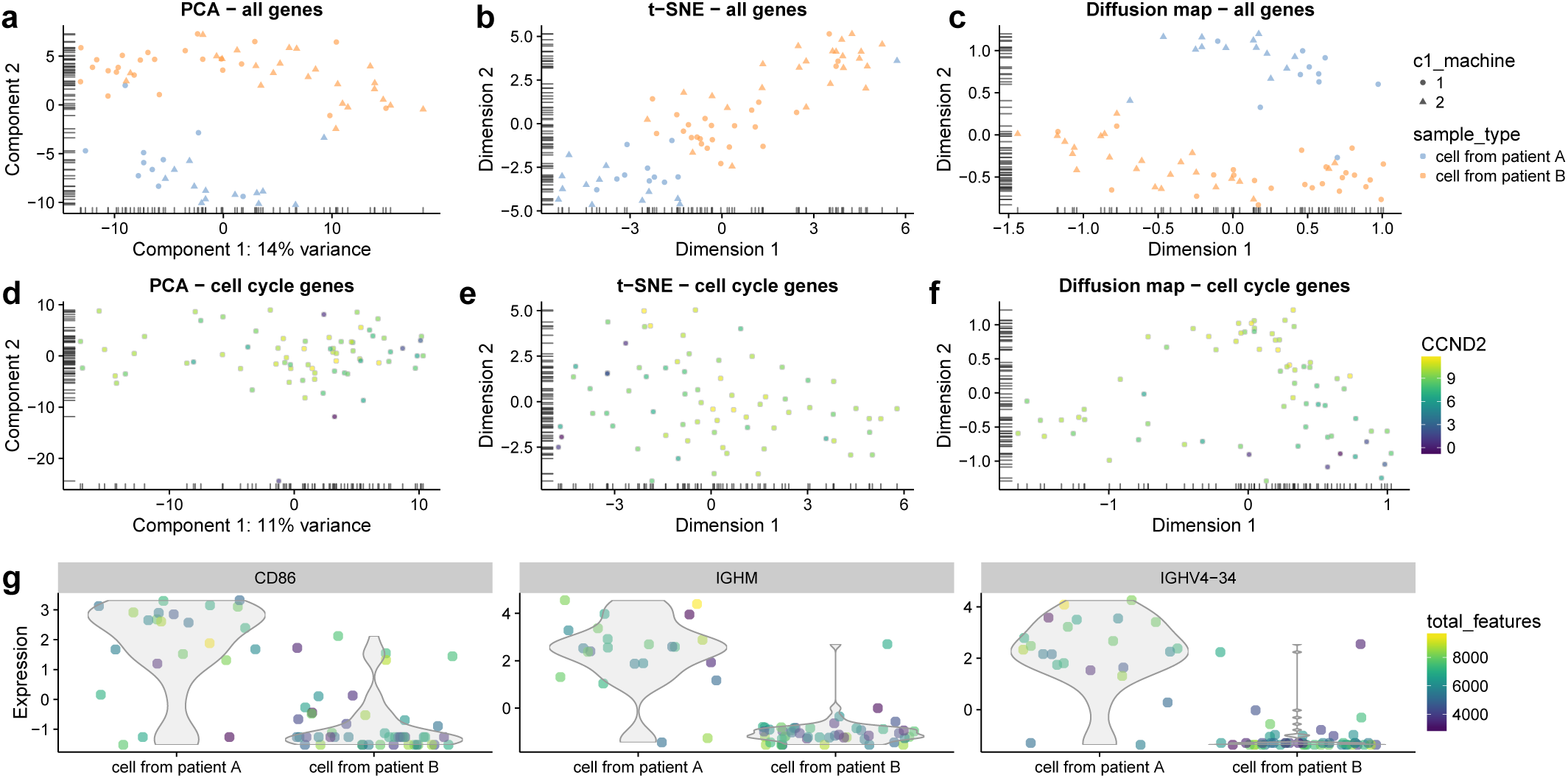
Reduced dimension representations of cells and gene expression plots with *scater*. Plots are shown using all genes (**a-c**) and cell cycle genes only (**d-f**) using PCA (**a,d**), t-SNE (**b,e**) and diffusion maps (**c,f**), where each point represents a cell. In the top row (**a-c**), points are coloured by patient of origin, sized by total features (number of genes with detectable expression) and the shape indicates the C1 machine used to process the cells. In the second row (**d-f**), points are coloured by the expression of *CCND2* (a gene associated with the G1/S phase transition of the cell cycle) in each cell. With the plotExpression function, gene expression can be plotted against any cell metadata variables or the expression of another gene—here, expression for the CD86, IGH44 and IGHV4-34 genes in each cell is plotted against the patient of origin (**g**). The function automatically detects whether the x-axis variable is categorical or continuous and plots the data accordingly, with x-axis values “jittered” to avoid excessive overplotting of points with the same x coordinate.

By default, the PCA and t-SNE plots are produced using the features with the most variable expression across all cells. We focus on the most variable genes to highlight any heterogeneity in the data that might be driving interesting differences between cells. Alternatively, we can apply *a priori* knowledge to define a set of genes that are associated with a biological process of interest, and construct plots using only these features. For example, Scialdone et al. (2015) found that using prior knowledge to define feature sets is vital for exploring processes like the cell cycle, which can have substantial effects on single-cell expression measurements (Buettner et al., 2015). The subsetting and filtering methods for SCESet objects facilitate the generation of reduced-dimension plots for particular gene sets, in order to investigate certain effects in the data such as those due to the cell cycle (Figure 3d–f).

The various types of reduced-dimension plots can be used to examine the structure of the cell population, including the formation of distinct subpopulations or the presence of continuous trajectories. Cell-level variables stored in the SCESet object can be used to define the shape, colour and size of points plotted, allowing more information to be conveniently incorporated into each plot (e.g., cells are coloured by *CCND2* expression in Figure 3d–f). The plotExpression function is also provided for plotting expression levels of a particular gene against any of the cell phenotype variables or the expression level of another feature (Figure 3g). This allows the user to inspect the expression levels of a feature or set of features in full detail, rather than relying only on summary information and reduced-dimension plots where information is necessarily lost.

### Data normalisation and batch correction

Scaling normalisation is typically required in RNA-seq data analysis to remove biases caused by differences in sequencing depth, capture efficiency or composition effects between samples. Frequently used methods for scaling normalization include the trimmed mean of M-values (Robinson and Oshlack, 2010), relative log-expression (Anders and Huber, 2010) and upper-quartile methods (Bullard et al., 2010), all of which are available for use in *scater*. In addition, *scater* is tightly integrated with the *scran* package that implements a method utilising cell pooling and deconvolution to compute size factors better suited to scRNA-seq data (Lun et al., 2016).

After scaling normalisation, further correction is typically required to ameliorate or remove batch effects. For example, in the case study dataset, cells from two patients were each processed on two C1 machines. Although C1 machine is not one of the most important explanatory variables on a per-gene level (Figure 2e), this factor is correlated with the first principal component of the log-expression data (Figure 2f). This effect cannot be removed by scaling normalisation methods, which target cell-specific biases and are not sufficient for removing large-scale batch effects that vary on a gene-by-gene basis (Figure 4a). Here we present two possibilities, all easily implemented in a scater workflow.

**Figure 4:**
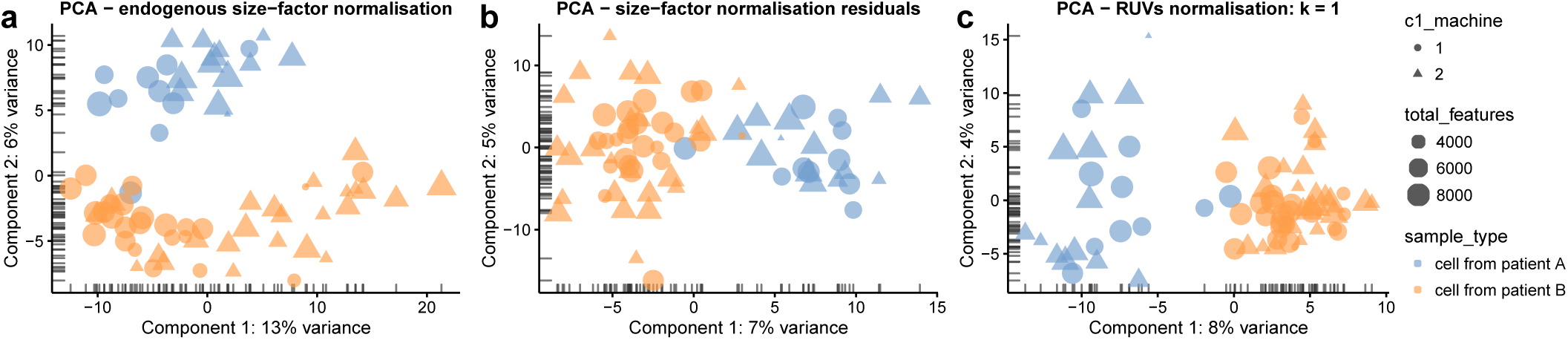
Normalisation and batch correction with *scater*. Principal component analysis plots showing cell structure in the first two PCA dimensions using various normalisation methods that can be easily applied in *scater*, including endogenous size-factor normalisation using methods from the *scran* package (**a**); expression residuals after applying size-factor normalisation and regressing out known, unwanted sources of variation (**b**); and removal of one hidden factor identified using the RUVs method from the *RUV* package (**c**). In all plots, the colour of points is determined by the patient from which cells were obtained, shape is determined by the C1 machine used to process the cells and size reflects the total number of genes with detectable expression in the cell.

The C1 machine effect is known from the design of the experiment, so we can easily regress out this effect in *scater*. With the normaliseExprs function the user can supply a design matrix of variables to regress out of the expression values, and residuals from the linear model fit can be used as expression values for downstream analyses. For the dataset here, we fit a linear model to the *scran* normalised log-expression values with the C1 machine as an explanatory factor. (We also use the log-total counts from endogenous genes, percentage of counts from the top 100 most highly-expressed genes and percentage of counts from control genes as additional covariates to control for these other unwanted technical effects.) We then use the residuals from the fitted model for further analyses (see Case Study in Supplementary Material). This approach successfully removes the C1 machine effect as a major source of variation between cells; the first principal component now separates the cells from the two patients, as expected (Figure 4b). This approach needs to be used carefully as single-cell data often deviate from normal distributions, but in many cases, as here, it can successfully ameliorate large-scale known batch effects.

In addition to removing known batch effects, it can be important for large data sets to identify (potentially unknown) sources of unwanted variation (Leek et al., 2010; Hicks et al., 2015; Grün and van Oudenaarden, 2015). *scater* is compatible with existing methods such as *svaseq* (Leek and Storey, 2007; Leek, 2014) and *RUVSeq* (Risso et al., 2014) to identify and remove these unwanted sources of variation, and the removeBatchEffect method in the *limma* package (Ritchie et al., 2015) to account for known batch effects. Here, just removing the first latent variable identified by the RUVs method from *RUVSeq* is sufficient to remove the machine effect, as the PCA plot now separates cells by patient rather than C1 machine (Figure 4c).

We emphasise that it is generally preferable to incorporate batch effects or latent variables into statistical models used for inference. Where this is not possible (e.g., for visualisation), directly regressing out these uninteresting factors is required to obtain “corrected” expression values for further analysis.

### Software and data integration

As part of the R/Bioconductor ecosystem, *scater* can be easily integrated with other software for scRNA-seq data analysis (Supplementary Figure 3). Because the SCESet class builds on existing Bioconductor data structures, most Bioconductor packages for expression analyses are able to operate seamlessly with SCESet objects. Tools that can integrate easily with *scater* include many options for data normalisation (Lun et al., 2016; Vallejos et al., 2015; Ding et al., 2015), differential expression analysis (Vallejos et al., 2016; Trapnell et al., 2014; Finak et al., 2015; Vu et al., 2016; Kharchenko et al., 2014; Korthauer et al., 2015; Andrews and Hemberg, 2016), heterogeneous gene expression analyses (Vallejos et al., 2015), clustering (Kiselev et al., 2016; Guo et al., 2015; Fan et al., 2016; Grün et al., 2015), latent or hidden variable analysis (Leek, 2014; Risso et al., 2014; Stegle et al., 2012; Chikina et al., 2015), cell cycle phase identification (Scialdone et al., 2015) and pseudotime computation (Trapnell et al., 2014; Angerer et al., 2015; Juliá et al., 2015; Campbell and Yau, 2016; Haghverdi et al., 2016). The *scater* package bridges the gap between raw reads and these downstream analysis tools by providing the pre-processing, QC, visualisation and normalisation methods and a data structure combining multiple data modalities and metadata necessary for convenient, robust and reproducible analyses of scRNA-seq data.

## Discussion

Single-cell RNA sequencing is widely used for high-resolution gene expression studies investigating the behaviour of individual cells. While scRNA-seq data can provide substantial biological insights, the complexity and noise of the data is also much greater than that of conventional bulk RNA-seq. Thus, rigorous analysis of scRNA-seq data requires careful quality control to remove low-quality cells and genes, as well as normalisation to adjust for biases and batch effects in the expression data. Failure to carry out these procedures correctly is likely to compromise the validity of all downstream analyses (Leek et al., 2010; Hicks et al., 2015; Grün and van Oudenaarden, 2015).

Here, we present an R/Bioconductor package, *scater*, that provides crucial infrastructure and methods for low-level scRNA-seq data analysis. The package introduces a data structure tailored to scRNA-seq data that is compatible with a vast number of existing tools in the Bioconductor project. The *scater* data structure combines multiple transformations of the expression data with cell and feature (gene or transcript) metadata and allows data sets to be easily standardised and shared. Wrapper functions for the popular RNA-seq quantification methods *kallisto* and *Salmon* facilitate the processing of raw read sequences to a SCESet object in R with expression data and accompanying metadata.

Quality control is a vital preliminary step for scRNA-seq and can be a time-consuming manual task. We present a tool for automated computation of QC metrics, novel plotting methods for QC and convenient subsetting and filtering methods to substantially simplify the process of filtering out unwanted or problematic cells and genes. The package provides a large array of sophisticated plotting functions so that cells can be visualised with a variety of popular dimensionality-reduction techniques in plots that incorporate cell metadata and expression values as plotting variables.

Normalisation is a critical aspect of scRNA-seq data processing that is supported by *scater*. Scaling normalisation methods, including the single-cell specific methods in the *scran* package, are seamlessly integrated into a *scater* workflow. Methods for identifying and removing batch effects and other types of unwanted variation are supported both with internal methods and through integration with a multitude of tools available in the R/Bioconductor environment. Once identified, important covariates and latent variables can be flagged for inclusion in downstream statistical models or their effects regressed out of normalised expression values.

Future development will include further extensions to data structures that will enable tight integration of single-cell transcriptomic, genetic and epigenetic data, as well as further refinement of the methods available as the single-cell field matures. Although *scater* has been produced for scRNA-seq data, its capabilities are well suited for single-cell qPCR data and bulk RNA-seq data, and may prove useful for supporting analyses of these data types.

## Conclusion

The *scater* package eases the burden for a user tasked with producing a high-quality single-cell expression dataset for downstream analysis. The intuitive GUI implemented in *scater* provides an easier entry point into rigorous analysis of scRNA-seq data for users without a computational background, enabling them to process raw reads into high-quality expression data within a single computing environment. Experienced users can take advantage of *scater*’s data structures, wide array of tools, suitability for scripted analyses and seamless integration with many other R/Bioconductor analysis tools. The data structures and methods in *scater* provide basic infrastructure upon which new scRNA-seq analysis tools can be developed. We anticipate that *scater* will be a useful resource for both analysts and software developers in the single-cell RNA sequencing field.

## Acknowledgements

We thank Marco Salvetti for provision of the two samples and Zam Cader for processing the samples. We would also like to acknowledge Peter Donnelly and Oliver Stegle for support and helpful discussions.

